# Heroin Type, Injecting Behavior, and HIV Transmission. A Simulation Model of HIV Incidence and Prevalence

**DOI:** 10.1101/591180

**Authors:** Georgiy V. Bobashev, Sarah Mars, Nicholas Murphy, Clinton Dreisbach, William Zule, Daniel Ciccarone

## Abstract

**Background and Aims:** Using mathematical modeling to illustrate and predict how different heroin source-forms: “black tar” (BTH) and powder heroin (PH) can affect HIV transmission in the context of contrasting injecting practices. By quantifying HIV risk by these two heroin source-types we show how each affects the incidence and prevalence of HIV over time. From 1997 to 2010 PH reaching the United States was manufactured overwhelmingly by Colombian suppliers and distributed in the eastern states of the United States. Recently Mexican cartels that supply the western U.S. states have started to produce PH too, replacing Colombian distribution to the east. This raises the possibility that BTH in the western U.S. may be replaced by PH in the future.

**Design:** We used an agent-based model to evaluate the impact of use of different heroin formulations in high- and low-risk injecting drug user populations who use different types of syringes (high vs. low dead space) and injecting practices. We obtained model parameters from peer-reviewed publications and ethnographic research.

**Results:** Heating of BTH, additional syringe rinsing, and subcutaneous injection can substantially decrease the risk of HIV transmission. Simulation analysis shows that HIV transmission risk may be strongly affected by the type of heroin used. We reproduced historic differences in HIV prevalence and incidence. The protective effect of BTH is much stronger in high-risk compared with low-risk populations. Simulation of future outbreaks show that when PH replaces BTH we expect a long-term overall increase in HIV prevalence. In a population of injectors with mixed low- and high-risk clusters we find that local HIV outbreaks can occur even when the overall prevalence and incidence are low. The results are dependent on evidence-supported assumptions.

**Conclusions:** The results support harm-reduction measures focused on a reduction in syringe sharing and promoting protective measures of syringe rinsing and drug solution heating.

## INTRODUCTION

In addition to being a huge burden on society in terms of drug overdoses and deaths, intertwined opioid and heroin epidemics in the United States (1, 2) have the potential to increase HIV transmission among people who inject drugs (PWID). Heroin use among American adults increased almost fivefold between 2002 and 2013 (3); most heroin is used by injection, and among PWID the proportion of persons living with a diagnosis of HIV infection in 2010 was 0.0215 (4). Among PWID, most HIV transmission is attributed to the sharing of needles and syringes (5); however, there is an intriguing difference in HIV prevalence among PWID in the western (5%-6%) versus eastern United States (11%-12%) (6, 7). A study of 96 U.S. cities showed that in the late 1990s and early 2000s the difference was 2%-11% versus 3%-35% with the mean in the western U.S. areas three times lower than that of the eastern states (3.6 vs. 9.4) (8). A similar relationship was observed in HIV incidence (9). In most of the western U.S. cities HIV incidence was less than 1 per 100 person-years, but the mean incidence in the eastern U.S. cities was over 3. Although a number of behavioral, environmental, historical, or structural factors may contribute to this disparity, we explore one plausible explanation based on the geographic distribution and use of specific heroin source-forms (6).

Heroin varies in physical and chemical characteristics by production source (10). To the west of the Mississippi River heroin is mostly available as a solid form (i.e., Mexican-sourced “black tar” [BTH]. “Powder heroin” (PH) is predominant east of the Mississippi River and although the source is changing from Colombian to Mexican, heroin *form* remains predominantly the same: powder (2, 11). Besides an ecological association between HIV prevalence, incidence, and geographic prevalence of BTH (Figure 1), these contrasting source-forms vary in physical state, cold/hot water solubility, pH, heat stability, weight/volume, and purity (6). These chemical features could provide mechanistic insight into the factors associated with differences in HIV incidence.

**Figure 1.**
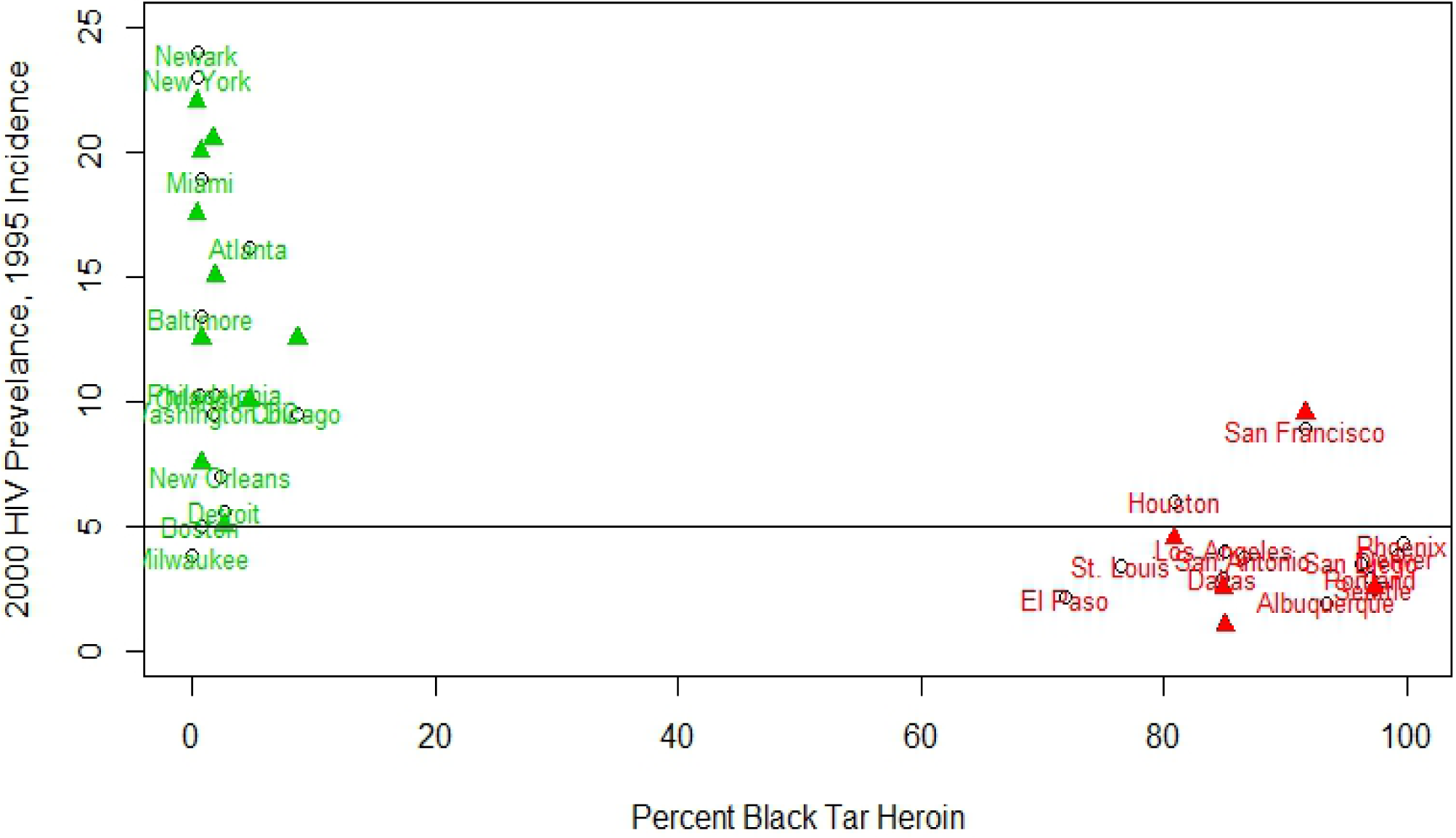
HIV prevalence and incidence in year 2000 vs. percentage of black tar heroin used in several major U.S. cities. Prevalence is depicted with city names. Triangles denote 1996 incidence per 500 person-years in cities where it was reported in Holmberg. (1996). Horizontal line represents 5% prevalence and incidence per 500 person years. Incidence was scaled to 500 person-years to get numeric values within the same range as the prevalence. HIV prevalence and incidence from Tempalski et al. (2009); prevalence of black tar in the cities as reported in Ciccarone and Bourgois, (2003) from Domestic Monitoring Program 1991–1993. Drug Enforcement Administration, U.S. Department of Justice.

### Heroin types, injecting practices, and HIV

HIV transmission can occur when a small amount of infected blood from a previous HIV-positive PWID remains in the syringe and enters the bloodstream of the next user. The striking difference in virus survivability when different source-types of heroin (BTH vs. PH) are used could substantially influence the probability of HIV transmission associated with heroin type (12). It has been thus hypothesized (6) that injection preparation practices associated with these heroin types explain the higher prevalence of HIV among PWID in cities with more prevalent PH versus more prevalent BTH. Despite some biological evidence that the virus does not survive well in high temperatures, actual survival and consequent HIV transmission are not well quantified for each individual injecting practice. In this study we attempted to identify and quantify behavioral and mechanistic factors affecting HIV transmission, and to translate them into HIV incidence and prevalence. Several mechanistic factors affect virus survivability (6):

1. Heat. BTH is usually heated to ensure better dissolution in water. Although some injectors also heat PH, it is done more out of tradition than out of need (6, 10).
2. Syringe rinsing. Because of its stickiness and viscosity, BTH requires multiple rinsing of the syringe to prevent the needle from clogging if it is to be reused (13, 14).
3. Switching from venous to muscular injection sites. BTH appears to induce faster and more severe venous scarring than PH, causing users to migrate to subcutaneous or intramuscular injection routes (15–17)
4. Acidity. Different heroin preparations vary widely by acidity (18). BTH is believed to be more acidic (pH ∼ 2.8) than PH (pH ∼ 4), with more acidic solutions showing reduced HIV survivability *in vitro* (19). However, the acidity of BTH has not been clearly quantified. These factors, combined with syringe-sharing practices and background HIV prevalence in the heroin-injecting community, affect the incidence and prevalence of HIV. Behavioral risk, incidence, and prevalence are related to each other in a dynamic way (i.e., risky behavior impacts future incidence, incidence and removal [e.g., mortality] rates impact temporal changes in prevalence). A dynamic simulation model that accounts for BTH/PH factors and risky behaviors can describe the emergence of the differences in HIV incidence and prevalence between U.S. geographic regions. In this study we developed such a model with two objectives in mind. One is to reproduce and quantify the historic evidence of the late 1990s and early 2000s described in Ciccarone & Bourgois (6). Another is to simulate potential “what if” scenarios. As Mexican-sourced heroin (both BTH and PH) increasingly dominates the heroin supply across the United States (11), one such scenario is Mexican-sourced PH completely replacing BTH. In the remaining sections of the manuscript we describe such a model and illustrate its application and findings.

## METHODS

In building a model we used the following mechanistic logic (Figure 2). A person can become HIV infected when a number of HIV viruses enters his or her bloodstream. This number (viral burden) is the product of the transmitted volume of liquid and viral load (concentration of virus in a unit of liquid). The use of BTH or PH could modify viral burdens (viral loads multiplied by the transmitted volume of liquid) and the volume of transmitted liquid can be altered by different injecting practices (e.g., high dead-space [HDS] syringes, sharing and rinsing). Thus, individuals with the same viral load in their bodies can transmit different amounts of virus depending on which heroin type they use and which injecting practices they employ. Injecting practices in turn can be affected by cultural realities such as injecting norms, the availability of replacement syringes, and the experience of the PWID involved. If one knows the combination of risk factors and the effect of each factor on HIV transmission, one can directly estimate the risk of HIV transmission. In reality, however, most of the evidence for the impact of each of these factors on HIV risk is either qualitative or indirectly quantitative; thus, we have to consider a number of simplifying assumptions based on available data.

**Figure 2.**
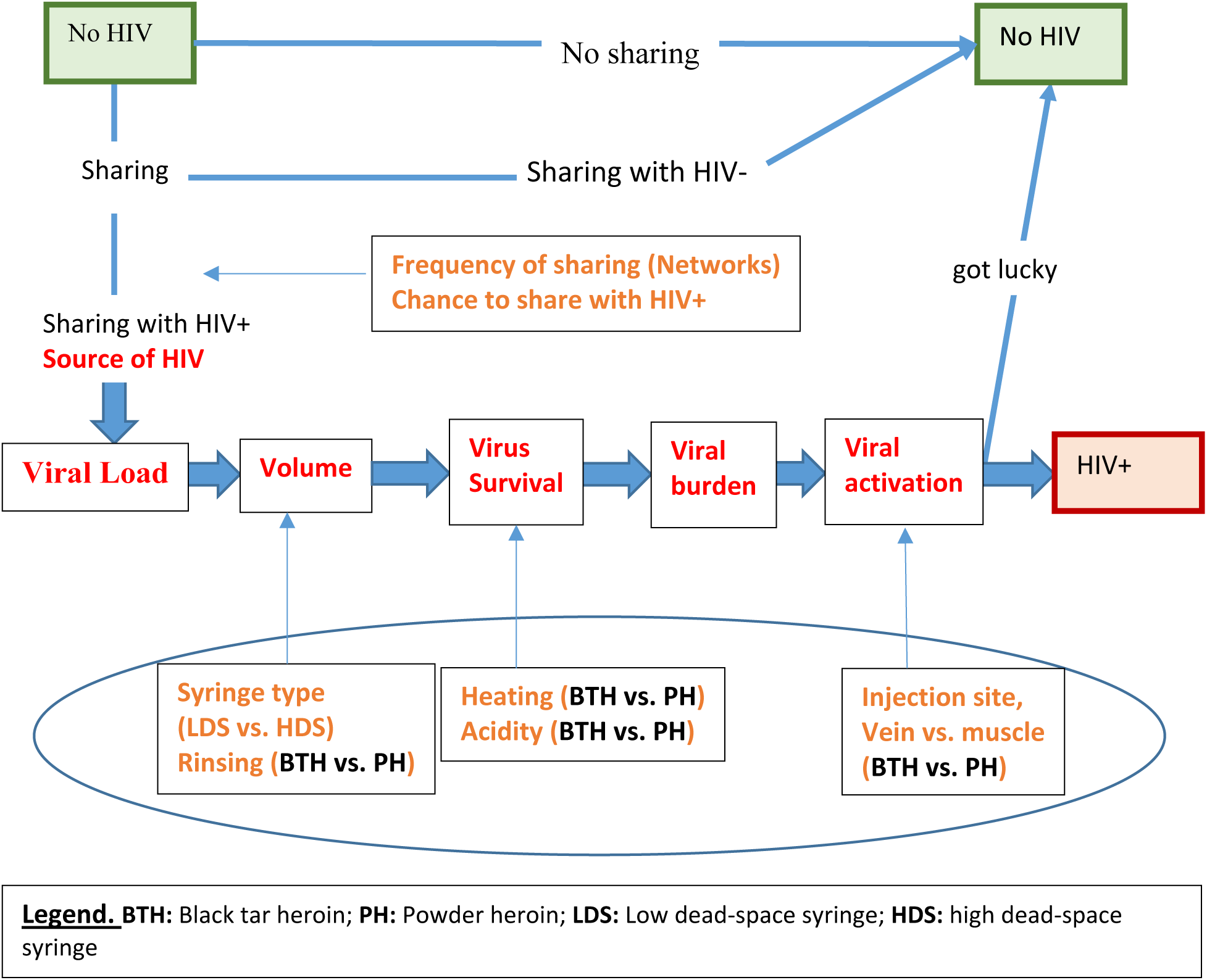
A depiction of factors affecting HIV transmission through sharing syringes and the role of BTH/PH in the probability of transmission. BTH/PH factors are overlaid with the oval.

In the following section we first describe an equation that links viral load and infectivity, then translate each of the heroin factors into the change in viral burden. Next, we describe a dynamic agent-based simulation model that combines all these factors together in the context of injecting behaviors and the structure of injecting networks.

### High and low dead space syringes

Zule et al. (20–22) and Bobashev and Zule (23) have shown that different syringe designs retain different amount of blood that can be transferred to the next injector during sharing. The syringes were classified as HDS or low dead space (LDS). Although the percentage of HDS syringes currently used by PWID in the United States is small (around 5%), in the late 1980s the percentage was higher and was critical to supporting HIV transmission, especially in high-risk populations that share syringes more than 10 times a year (23).

### Reference transmission probability per exposure

Estimates of the probability of HIV transmission per exposure through receptive syringe sharing range from 0.0051 to 0.0189, depending on the genetic subtype (B vs. E) (24–26). We thus use a reference value *P*_*ref*_ of 0.008, which is similar to the ones in Patel et al. (27) and Bobashev and Zule (23). Because most of the estimates of the HIV risk were obtained from studies in Thailand, we assume that users injected PH and used HDS syringes (20). We assume that most sharing occurred during the latent phase of HIV when the viral burden was around 10,000 copies per milliliter.

### Viral burden and HIV probability by a single virion

In modeling the relationship between the viral burden and probability of transmission we used a simple assumption that each virion has the same chance to start a disease. Thus, for a viral burden of *X* copies of virus the probability of starting a disease *P* will be

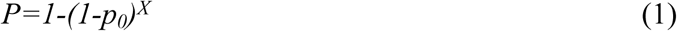

Which could be solved for *p*_*0*_ as

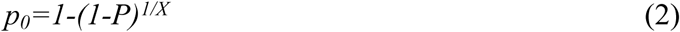

Where *P* is a probability of transmission associated with a viral burden *X* and *p*_*0*_ is a probability of starting an infection by a single virion. The estimates of *p*_*0*_ are difficult to obtain directly but could be evaluated from the following indirect considerations.

The volume of the shared blood and viral load is related to the type of syringe as described in Zule et al. (21, 22). If all shared syringes were rinsed once, the shared volume of blood retained in the dead space would be around 0.0063ml resulting in 63 copies of the virion. For a reference value of transmission probability *P*_*ref*_ = 0.008 equation (2) produces an estimate of *p*_*0*_ equal to 1.3*10^−4^. If all syringes are rinsed twice, the volume of shared blood is 0.001ml (10 copies of virion) and *p*_*0*_ equal to 8*10^−4^. We thus consider the estimate of *p*_*0*_ to be between 1.3*10^−4^ and 8*10^−4^.

### Heroin type factors and viral burden

#### Heating the solution

Following Clatts et al.’s (12) observations of cooking times and temperatures achieved in cookers and syringes in New York and Denver we considered that the solutions on average reach a temperature of 65°C (range 10-100) and it took about 6 seconds (range 3-10) to reach the mean temperature. In that study, when heated to 66°C in one sample the virus was not recoverable after 10 seconds and in another sample, HIV was not recoverable after 7 seconds when heated to 64°C. Heating for less than 5 seconds had some effect but not as strong as for heating over 5 seconds. It is notable, however, that extensive heating capable of eliminating the virus does not happen all the time. In Denver where BTH is predominantly used, about 20% of the time it was heated for less than 10 seconds and 40% of the time it did not reach 66°C. Thus, although protective, heating practices of BTH leave a window of opportunity for HIV transmission. We consider that when solutions of BTH are heated up to 66°C for over 10 seconds all viruses in the cooking equipment are inactivated. When heated liquid is drawn into syringes the temperature drops by about 2°C and the virus remaining in the syringe might still survive. We assume that in those 40% of heating events when the temperature does not reach 66°C, virus survival is about 70%. PH is either mixed cold or heated for up to 15 seconds, reaching 66°C for over 5 seconds only 10% of the time. Thus, on average, when heating the BTH solution, the survival rate of HIV virus in a syringe is around 10% (1-0.6*100%-0.4*70%), compared to 90% (1-10%) when using PH solution heated to lower temperature.

#### Rinsing syringes

Most PWID rinse their syringes at least once if they expect to use them in the future to prevent blood from clotting and clogging the needles. However, ethnographic observations show that among BTH users an extra syringe rinse became the norm to additionally ensure that tar residuals do not clog future injections. Following (23, 28) we assume that 0.084ml of liquid is left in an HDS syringe and that a 1ml syringe is rinsed with 0.5 ml of water. Such a rinse reduces the amount of blood (and thus a number of virions) by a factor of 0.084/0.5 = 0.17. For LDS the reduction is much stronger and is around 0.002/0.5 = 0.004.

#### Acidity

In vitro experiments show that HIV-contaminated syringes are significantly less likely to yield recoverable HIV when rinsed with citric acid solutions of pH at or less than 2.3 (19). Although there is measured evidence of highly acidic solutions (depending on the type of acidifier) in Europe (18), data on the acidity of BTH solutions is poorly known, although suspected to be acidic (is produced with acetic acid rather than acetic anhydride used in PH (29). We thus do not use any specific HIV survival factor associated with acidity but note that our estimates are conservative (i.e., the effect of BTH on killing the virus is likely stronger because of the additional effect of acidity).

#### Switching to non-intravenous injection sites

Using data from accidental needle-stick injuries, the risk of HIV transmission following intramuscular poke is about 3 times less compared to intravenous poke (0.002 vs. 0.006) (30). We assume from ethnographic observations that on average about 25% of BTH injections are intramuscular or subcutaneous (32). Rich et al. (17) suggest that the probability of HIV transmission through subcutaneous injection is negligible.

### Simulation model

Agent-based models are microsimulations that reproduce individual behaviors in the context of social networks and account for environmental and individual risks. In our model we simulate communities of PWID where individuals form injecting networks, can share syringes and equipment, and transmit HIV from one person to another. These models allow detailed description of injecting practices where HIV transmission risk is detangled into individual components: sharing LDS syringes, sharing HDS syringes, and using PH versus BTH. Injecting networks in our model have different sizes and connectivity to imitate urban and rural areas and different injecting norms (high and low probability of sharing). The worst-case scenario is set up to consider a highly dense network with high-risk behaviors of agents who use a large proportion of HDS syringes and inject PH. On the other extreme, best case scenario, we consider a network with low level of sharing, use of LDS syringes, and BTH. The latter scenario leads to very low HIV incidence and illustrates an exemplary success case of harm reduction. All other scenarios are between these two extremes. One such scenario is when individuals switch from BTH to PH and less experienced individuals join the network without knowledge of or adherence to the safe norms. Specific model setting assumptions are described below.

- The model focuses on the injecting pathway of HIV transmission, and sexual transmission is considered only as a background force of infection, so each individual can become infected with some small per act probability (0.001) scaled down by overall HIV prevalence (probability that sexual partner is HIV positive) and assuming a frequency of sexual activity of once per week.
- In our model the agents were arranged in clustered networks (i.e., more intense syringe sharing with the members of the same networks [buddies] and only occasional sharing with the members of other networks [strangers]). These strangers represent the common pool of injectors and facilitate the spread of HIV from one network to another. In our model the structure of networks (i.e., who injects with whom) and within-network risk behavior (i.e., how often agents share the syringe) was drawn at random from uniform distributions. These networks were of different sizes: 20% very small (2 individuals), 70% medium (3-8 individuals), and 10% large (9-20 individuals) with the average network size of eight as was observed in a study of PWID in North Carolina. We considered for simplicity 64 networks that resulted in approximately 500 PWID.
- Each network could exhibit either high- or low-risk behavior norms. High versus low risk is defined as higher rates of syringe-sharing episodes with strangers (1 vs. 12 times a year) and within a network (e.g., 5 vs. 30 times a month). These parameters were estimated from the supplementary data collected in two studies of PWID (28, 33).
- The role of different types of syringes and sharing practices in HIV transmission has been discussed in a number of papers (23, 28, 33–35). Most of the current injecting in the United States occurs with LDS; however, for simulations related to the period 1992-2002 we assume that about 10%-20% of syringes used by PWID were HDS, with higher rates in the south and lower in the west (22, 33).
- We assume that only a fraction (e.g., 0.5) of the individuals in a network is present at a particular injecting episode. In simulation we only track injections that result in sharing syringes and do not consider injections alone.
- We consider a removal rate of 4%. Individuals may be removed because they die or stop injecting. To keep the population stable and have the same denominator for incidence and prevalence calculations, removed individuals are replaced with a similar but HIV-negative individual on the basis that most new injectors are unlikely to have been exposed to the virus.
- Following Jaquez et al. (36), and Fiebig et al. (37) we assume that during the acute stage of HIV infection the probability of transmission increases between 5 and 30 times. For simplicity we use the value of a 10-fold increase.
- Our model does not consider a number of factors that affect the dynamics of HIV spread, such as behavior change after becoming HIV positive, number of times syringes are reused, and variation in viral load resulting from the use of antiretroviral therapy. These factors affect the actual incidence and prevalence; however, for the purpose of examining the effects of heroin types, we assume that these factors vary in the same way between users of PH and BTH, and the additional parameterization will increase model complexity without critically impacting the results. A list of parameters is presented in Table 1.

The model was programmed in NetLogo to provide interactive visualization of HIV transmission in a community. A screenshot of the user interface is presented in Figure 3.

**Table 1.** Model Parameters and Their Sources*.

| **Model Parameter** | **Value**<br>**High risk/Low risk** | **Source** |
| --- | --- | --- |
| Initial HIV prevalence | 0.05 | Experimental parameter |
| Number of networks | 64 | Experimental parameter |
| Size of network cluster | 2–17, average 8 | Bobashev and Zule, (2010) |
| Proportion of people in the cluster participating in sharing | 0.5 | Experimental parameter |
| Number of times sharing per month with buddies | 30/5 | Zule et al. (2002, 2009) |
| Number of times injecting with a stranger | 10/1 per year | Experimental parameter |
| Removal rate (includes PWID HIV+ all-cause mortality and PWID leaving the population [i.e., stop injecting]). | 0.04 per year | Bailey et al. (2007), De et al. (2007) |
| Risk of HIV transmission assuming HDS and PH | 0.008 | Hudgens et al. (2001, 2002) |
| Risk multiplier for an acute stage of HIV infection | 10 | Jacquez et al., Fiebig et al. (range 5-30) |
| Risk multiplier for muscle injection | 0.29 | Bagaley et al. (2006) |
| Risk multiplier for heating | 0.1 | Clatts et al. (1999) |
| Risk multiplier for an extra rinse of an HDS syringe | 0.17 | Zule et al. (1997, 2018) |
| Risk multiplier for an extra rinse of an LDS syringe | 0.004 | Zule et al. (1997, 2018) |
| Annual chance of sexual HIV | 10-4 | CDC (2017) incidence rates through sexual contacts |

Note. SATH-CAP = Sexual Acquisition and Transmission of HIV Cooperative Agreement Program; PWID = injecting drug user; BTH = black tar heroin, PH = powder heroin, HDS = high dead-space syringe; LDS = low-dead-space syringe.
\*“Experimental” parameter means that the value of the parameter varies between different geographic areas, cultures, etc. For illustration purposes, we use a value that is considered reasonable. We use experimental parameters to design “what if experiments” aimed at evaluating intervention strategies and conducting sensitivity analysis.

**Figure 3.**
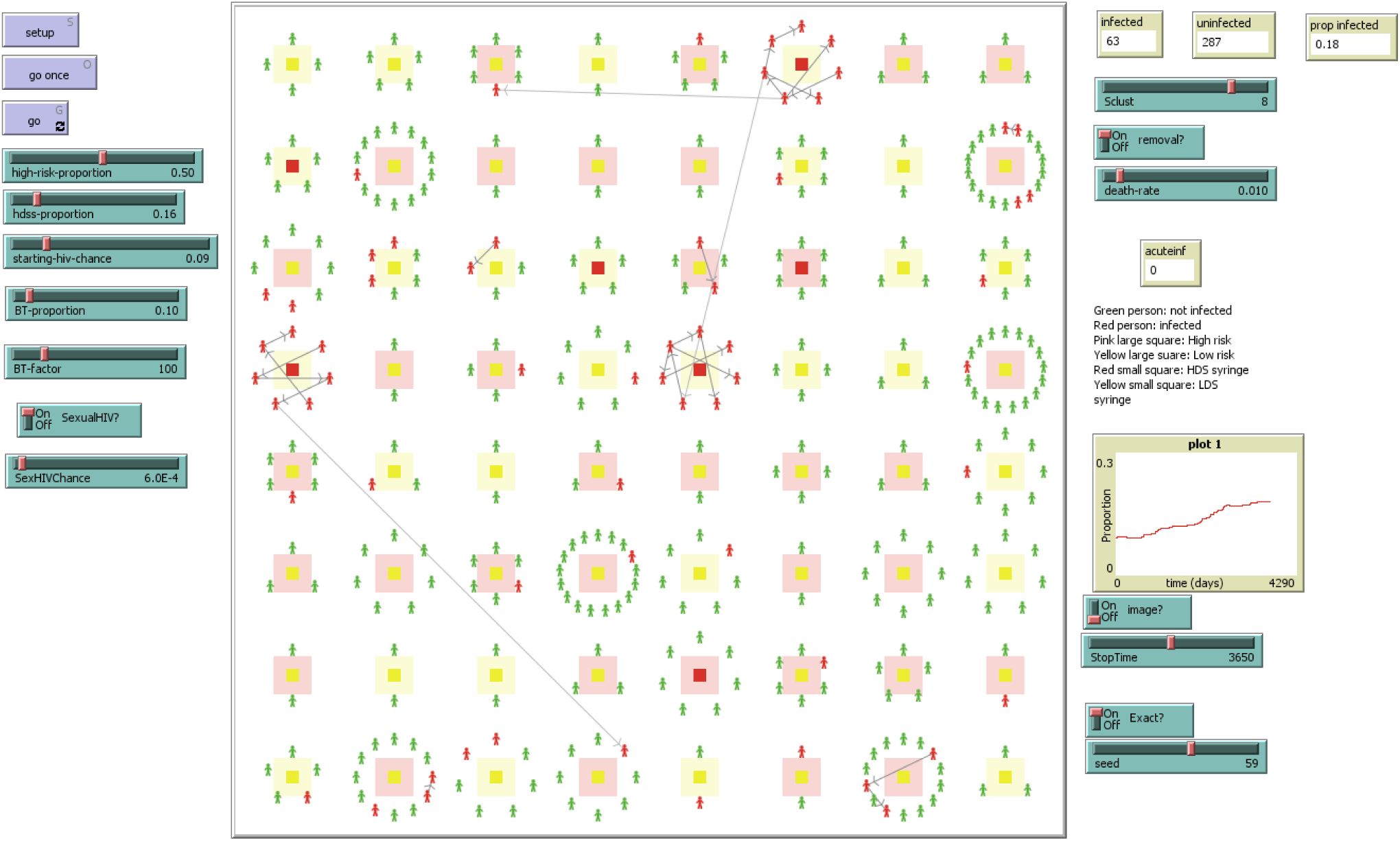
A screenshot copy of the simulation model that describes the spread of HIV among PWID. Each injecting network can be high or low risk in terms of frequency of syringe sharing (large pink or yellow squares) and using HDS or LDS syringes (small pink or yellow squares inside the large squares).

### Simulation scenarios

In our analysis we considered the following scenarios:

#### Historic scenarios (Scenario A1, A2)

These scenarios simulate what could potentially be the historic setting east and west of the Mississippi River leading to observations described in Ciccarone and Bourgois 2003 (6):

Scenario A1. BTH + 20% HDS. This scenario mimics the western U.S. around the late 1980s through late 1990s. We assume that HDS syringes were used about 20% of the time even with the early introduction of needle and syringe programs. We assume 100% of double rinses and a starting HIV prevalence of 3%.

Scenario A2. PH + 20% HDS. This scenario mimics the eastern U.S. around the late 1980s through late 1990s. Here we assume 60% chance of the second rinse and a starting HIV prevalence of 4%.

Futuristic scenario (Scenario B). A hypothetical scenario where PH becomes the dominant form of heroin on the market across the United States, almost completely eliminating the use of BTH.

Scenario B. PH + 97% LDS. This scenario assumes that LDS syringes will be used almost exclusively to inject heroin, but occasionally HDS syringes are used. This could be true especially in areas where needle and syringe programs have limited reach.

We computed incidence and prevalence trajectories for the simulated PWID population and for high- and low-risk individuals under different combinations of parameter values. We repeated simulations 1,000 times to obtain stable and smooth estimates of HIV prevalence and incidence and uncertainty bounds. At each simulation we considered uncertainty in parameter values (by drawing a value from a distribution), structural uncertainty (the number of networks and their structure), and internal stochasticity (randomness in behavior and disease transmission).

## FINDINGS

### The role of BTH from a historical perspective

By considering communities with both 95% and 5% BTH use, our model was able to simulate HIV prevalence among PWID in the western (3%-10%) and eastern United States (3%-20%), resulting in HIV rates similar to those observed in Holmberg et al. (9) and Tempalski et al. (8). In Figure 4 we present simulated HIV prevalence in the eastern and western United States for different proportions of high-risk networks. Simulated prevalence depends on the proportions of high-risk networks, but that dependence is small in the areas with a high proportion of BTH but quite substantial in the areas with PH.

**Figure 4.**
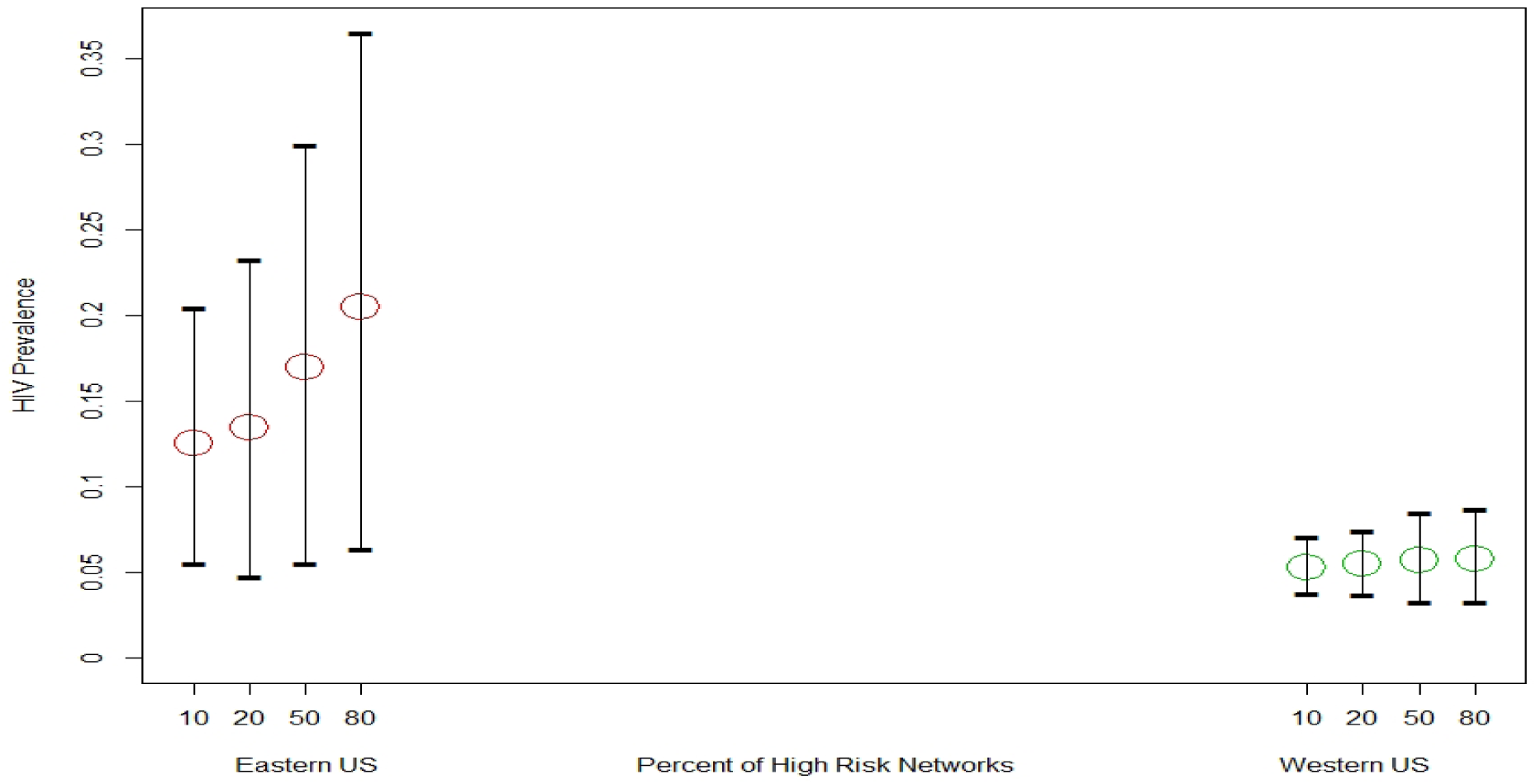
Simulated HIV prevalence in eastern cities (5% BTH) and western cities (95% BTH) for different percent (10%, 20%, 50%, 80%) of high-risk networks. Bars denote 95% range of simulated prevalences.

### The role of PH from the perspective of dominance in the illicit drug market

Simulation results show a significant increase in the incidence and prevalence of HIV if PH replaces BTH in a community. When simulating what could happen in the western United States if PH became dominant there, we started with an average HIV prevalence of 5% and 20% of high-risk networks. In 10 years, the HIV prevalence increased to 8%, but the variability of outcomes was high ranging from 4% to 17%. For high proportion (0.8) of high-risk networks the mean prevalence became 13% but the variability of the outcomes was even higher ranging from 4% to 30% as illustrated in Figure 5.

**Figure 5.**
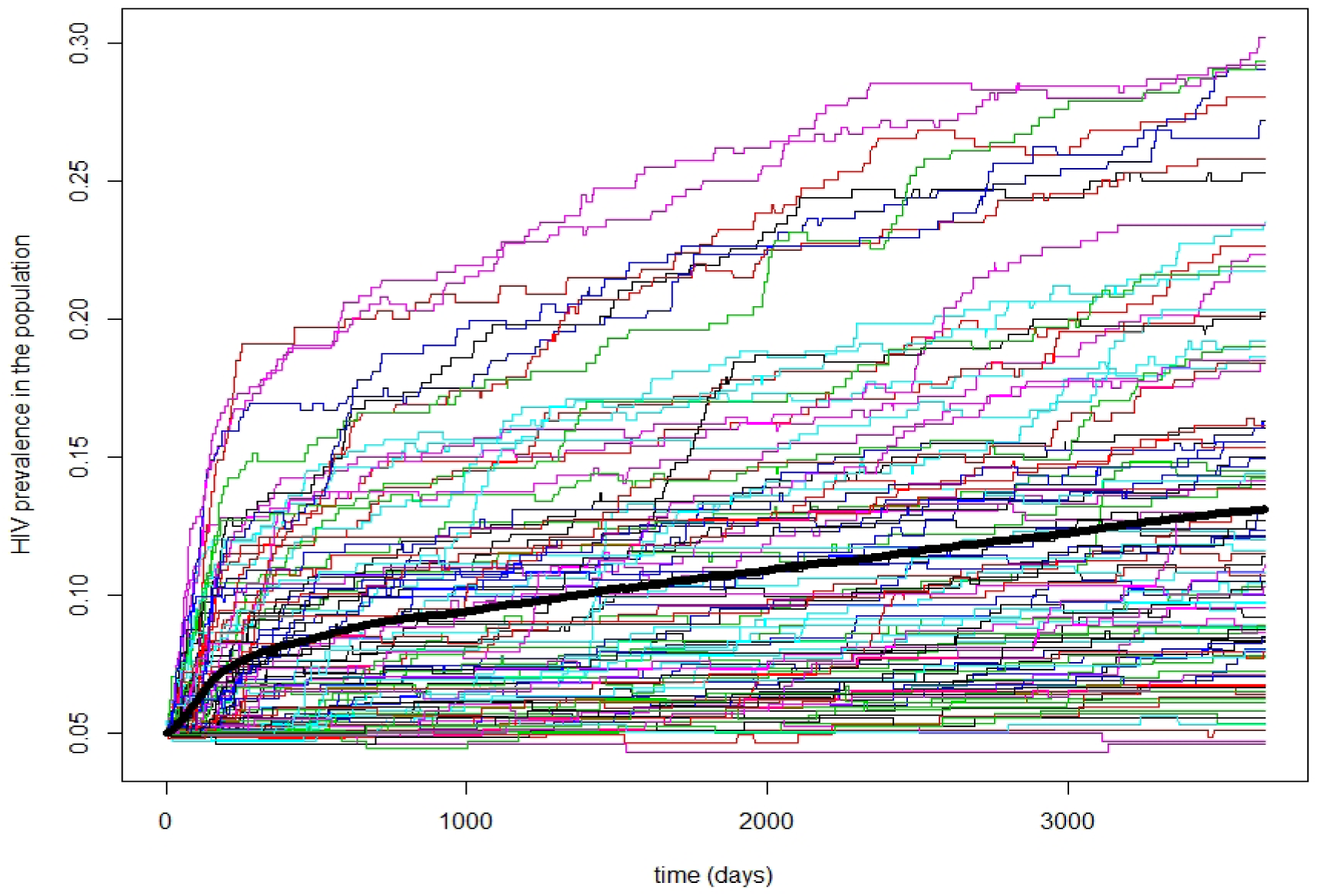
An example of high variability of 100 possible HIV trajectories when a community of PWID with 80% of high-risk networks switches to PH. The mean trajectory is shown in a thick solid black line. Variability of trajectories around the mean is high with a few trajectories (8 in this example) reaching a prevalence of over 25%. When HIV reaches a large high-risk cluster, the trajectory shows a quickly rising increase in prevalence.

## DISCUSSION

We have presented an agent-based simulation model that explores the effects of BTH and PH on HIV prevalence and provides a plausible explanation for the differences in HIV prevalence between metropolitan areas in the eastern and western United States. We have also shown that if PH replaces BTH, the overall prevalence of HIV will slowly grow, primarily because there will be local HIV outbreaks. These outbreaks are driven by local high-risk networks. Although many experienced users over time have accepted certain safer injecting norms, newer, less experienced users (e.g., migrant groups or users who are transitioning to heroin use from prescription opioids) might be at the highest risk. The role of viral load, especially in the acute phase of HIV, becomes additional fuel in high-risk networks with a lack of harm reduction norms.

New populations of users who are joining high-risk networks might not be exposed to harm reduction norms and will thus suffer. Because syringe exchange programs have been shown to substantially reduce syringe sharing (38, 39) our simulations implicitly suggest that new pockets of HIV could emerge in areas where needle and syringe programs are lacking and PWID routinely share their equipment. According to the Foundation for AIDS Research (40) currently needle and syringe exchange programs remain illegal in 15 states (MS, AL, GA, SC, FL in the South and TX, OK, KS, NE, SD, IA, MO, AR, WY, ID in the West). Even in states where needle and syringe programs are legal, rural areas often lack access to them, and 93% of counties that are under high risk of HIV/HCV outbreak do not have these programs (40). The opioid crisis has brought some pill users to injection practices, whether by transitioning to injecting heroin (41) or through injecting opioid pills, opening a specific opportunity for HIV outbreaks. In Scott County, Indiana, between 2014 and 2015, 181 people were diagnosed with HIV infection in a county that had seen only 5 diagnoses from 2004 to 2013. Eighty-eight of those diagnosed reported injection use of extended-release oxymorphone which may be crushed, dissolved, and cooked (42, 43). Our simulations underscore that where the opioid/heroin epidemic is moving, so should be moving interventions such as harm reduction and robust needle and syringe programs. Although the initial motivation for our paper was the role of BTH in the early HIV epidemic, our conclusions have a much broader contemporary application internationally, for example in Tajikistan and other countries where PH is used, needle and syringe programs routinely distribute HDS syringes and injecting norms (e.g., heating) are not preventing viral transmission (21, 35).

### Sensitivity analysis and uncertainty assessment

The simulation model shows that the results are naturally sensitive to the rates of sharing and risky behavior factors. We observed a strong interaction between behavioral risk level, effects of BTH, and use of HDS syringes. In general, the higher the risk, the more other protective factors (BTH or LDS) have an effect. This leads to an important observation regarding public health interventions. The effect of BTH is strong in high-risk networks (i.e., networks where agents share often within their network and with agents from the other networks [strangers]). For low-risk networks the protective effect of BTH is negligible as seen in Figure 4. These results offer a simple explanation that when a network is of low risk (an extreme situation is when no sharing of any equipment occurs) then HIV is not spreading through injecting contacts regardless of whether the network is using PH or BTH. When the level of sharing is high and is more aggravated by unsafe sharing practices the protective role of BTH becomes more evident.

Figure 5 illustrates the variability of the trajectories for a single scenario. Additional variability of network structures, between-person variability in viral load, and structural uncertainty (i.e., different modelers would model and program the same process differently) will increase the spread of possible outcomes. Adding these levels of uncertainty will likely result in coverage of the entire range of possibilities (i.e., in some simulations some runs will produce prevalence of 1). Although such outcomes are very unlikely, for small communities such as Scott County, IN, high-risk behavior could lead to very high incidence and then prevalence.

### Limitations and data considerations

As with most simulation studies, ours has a number of common limitations. The utility of a simplified simulation modeling is limited because the results are dependent on a number of simplifying assumptions. Some assumptions are technical (e.g., the structure of the injecting networks, homogeneous mixing within networks). Others are critical to the description and understanding of the underlying process. For example, as is mentioned in Tempalski et al. (8) there are many explanations for why HIV prevalence and incidence among PWID is so different between U.S. regions. They mention “program efforts to increase users’ access to clean syringes both through syringe exchange programs and pharmacies; efforts to promote safer injection practices; effects of antiretroviral therapies on infectivity of PWID, deaths from HIV not being matched by new infections; and possible changes in risk networks and other social mixing patterns which vary from place to place. Differences in HIV prevalence rates may also reflect differences in availability, accessibility and effectiveness of HIV prevention and treatment programs across metropolitan areas.” More explanations can be found in Ciccarone and Bourgois (6). These are all important factors and could be further considered in the models. For our simple exercise we rolled up many effects into a few proxy factors. For example, access to clean syringes and effectiveness of prevention practices are rolled into mechanistic risky behavior factors related to HIV transmission (i.e., how often PWID share syringes with “buddies” and “strangers”). Access to HIV treatment programs could be also represented by the level of risky behavior and the level of HIV viral load, which could then be translated into the probability of transmission.

One critical limitation in our analysis is that we do not consider injections of cocaine, crack and methamphetamine. A certain portion of HIV transmission among PWID occurs among this population but the data to calibrate the difference between injecting only heroin, heroin and other substances, and other substances but not heroin is problematic and is beyond the scope of this study.

In our model, we specifically focused on a question that relates BTH and PH availability and the role the related injecting practices can have in HIV transmission when keeping all other factors the same. From this perspective, modeling studies allow an unparalleled opportunity to explore “what if” scenarios that are not possible in the real world. Although simulation results do not necessarily reproduce real-life processes, they do provide insight into the variability of HIV prevalence as a function of several additional risk factors, such as frequency of syringe sharing, mixing patterns, and specific use practices. Another factor that justifies such simplification is the lack of reliable data about these practices. Ethnographic research provides in-depth insights into specific behavior, but it is very localized, while epidemiological studies are broad but lack depth. Some measurements are just not ethical and not feasible to obtain. For example, ethical concerns prohibit using controlled human experiments to estimate the dependence of HIV transmission on viral load.

Because the aim of this research is to show the *relative* effect of different types of heroin, the model is simple and focused to illustrate the main point. An interesting, emerging new source of data is online blogs and help sites that can be mined for useful information. This information provides complementary insights into what users (and not researchers) think is important. Our future research will explore these new avenues.

## Acknowledgements

This study was funded in part by the National Institutes of Health grant R01 DA037820, Ciccarone PI

